# Characterizing the dynamic and functional DNA methylation landscape in the developing human cortex

**DOI:** 10.1101/823781

**Authors:** Kira A. Perzel Mandell, Amanda J. Price, Richard Wilton, Leonardo Collado-Torres, Ran Tao, Nicholas J. Eagles, Alexander S. Szalay, Thomas M. Hyde, Daniel R. Weinberger, Joel E. Kleinman, Andrew E. Jaffe

## Abstract

DNA methylation (DNAm) is a key epigenetic regulator of gene expression across development. The developing prenatal brain is a highly dynamic tissue, but our understanding of key drivers of epigenetic variability across development is limited. We therefore assessed genomic methylation at over 39 million sites in the prenatal cortex using whole genome bisulfite sequencing and found loci and regions in which methylation levels are dynamic across development. We saw that DNAm at these loci was associated with nearby gene expression and enriched for enhancer chromatin states in prenatal brain tissue. Additionally, these loci were enriched for genes associated with psychiatric disorders and genes involved with neurogenesis. We also found autosomal differences in DNAm between the sexes during prenatal development, though these have less clear functional consequences. We lastly confirmed that the dynamic methylation at this critical period is specifically CpG methylation, with very low levels of CpH methylation. Our findings provide detailed insight into prenatal brain development as well as clues to the pathogenesis of psychiatric traits seen later in life.

## Introduction

DNA methylation (DNAm) plays an important role in the epigenetic regulation of gene expression, tissue differentiation, and development from fetal life through adolescence and likely beyond. It has been shown that in the human brain, DNAm is particularly plastic during the first five years of postnatal life, both at CpG and CpH (H=A, T, or C) sites ^1^. Methylation levels at key sites change over time and these changes lead to adjustments in gene expression and splicing. These key regions have also been linked to neurodevelopmental disorders such as schizophrenia, in which early dysregulation plays a vital role ^2,3^. DNAm is an attractive epigenetic mechanism to study in post-mortem human brain tissue, because it represents an interaction between genetic and environmental effects. External factors such as changes in diet ^4^, exposure to cigarette smoking ^5^, and exposure to arsenic ^6^ have been associated with both global and site-specific changes to DNAm levels.

In order to better understand the causes and consequences of deviant DNAm patterns in psychiatric disease development, we must first understand the normal landscape. Illuminating typical methylation changes in prenatal development will both provide insight into gene expression and molecular pathways active in the postnatal developing brain, and provide a baseline for identification of aberrant DNAm in postnatal disease states, as the pathological changes that lead to symptoms of psychiatric disease may precede the onset of illness by several decades ^7^.

The dorsolateral prefrontal cortex (DLPFC) is a dynamic region of the brain throughout development, essential for motor planning, conceptual organization, and working memory, and is often functionally dysregulated in patients with schizophrenia ^8^. Previous studies using microarrays to quantify DNAm have revealed that there are many DNAm changes in the DLPFC around the time of birth ^9^, but the dynamics of the prenatal brain are far less characterized.

Previous work assessing fetal brain DNAm has largely used microarrays, and sampled whole brain rather than a specific region ^10^. Most cells in the developing brain are neuronal ^11^, and it has been shown that neuronal DNA methylation is especially dynamic in the earliest stages of postnatal life, making the DLPFC a potentially fruitful region for deeper interrogation in prenatal samples ^1^. In the present report, we describe the use of whole genome bisulfite sequencing (WGBS) to capture an unbiased map of the DNAm landscape, and to characterize both CpG and CpH methylation during prenatal brain development. Dynamic regions of DNAm even in the early developmental context are likely to be associated with psychiatric risk-associated genes, and connection to gene expression data can validate the importance of these genomic regions. These important regions will help point to pathways and mechanisms of normal brain development as well as psychiatric and neurodevelopmental conditions.

## Materials and Methods

### Study samples

Brain tissue from these second-trimester prenatal samples was obtained via a Material Transfer Agreement with the National Institute of Child Health and Human Development Brain and Tissue Bank. All specimens were flash-frozen, then brain pH was measured and postmortem interval (PMI, in hours) was calculated for every sample. Postmortem tissue homogenates of the dorsolateral prefrontal cortex (DLPFC) were obtained from all subjects. Samples were obtained from the developing prefrontal cortex from the dorsolateral convexity of the frontal lobe half-way between the frontal pole and temporal pole, 4 mm lateral to the central sulcus. Specimens extended from the surface of the brain to the ventricular zone. There were 10 of each male and female subjects, and 17 subjects were African American and 3 were European ancestries (Table S1).

### Data generation

Genomic DNA was extracted from 100 mg of pulverized DLPFC tissue with the phenol-chloroform method. DNA was subjected to bisulfite conversion followed by sequencing library preparation using the TruSeq DNA methylation kit from Illumina. Lambda DNA was spiked in prior to bisulfite conversion to assess its rate, and we used 20% PhiX to better calibrate Illumina base calling on these lower complexity libraries. Resulting libraries were pooled and sequenced on an Illumina HiSeq X Ten sequencer with paired end 150bp reads (2x150bp), targeting 90Gb per sample. This corresponds to 30x coverage of the human genome as extra reads were generated to account for the addition of PhiX.

### Data Processing

The raw WGBS data was processed using FastQC to control for quality of reads ^12^, Trim Galore to trim reads and remove adapter content ^13^, Arioc for alignment to the GRCh38.p12 genome (obtained from ftp://ftp.ncbi.nlm.nih.gov/genomes/all/GCA/000/001/405/GCA_000001405.27_GRCh38.p12/GCA_000001405.27_GRCh38.p12_assembly_structure/Primary_Assembly/assembled_chromosomes/) ^14^, duplicate alignments were removed with SAMBLASTER ^15^, and the Bismark methylation extractor to extract methylation data from the sequencing data ^16^. We then used the bsseq R/Bioconductor package (v1.18) to process and combine the DNA methylation proportions across the samples for all further manipulation and analysis ^17^. After initial data metrics were calculated, the methylation data was smoothed using BSmooth for modelling. CpGs were filtered to those that had ≥ 3 coverage in all samples, and CpHs were filtered to those that had ≥ 3 coverage and non-zero methylation in at least half (≥10) of the samples.

### Comparison to 450k

In order to compare WGBS methylation levels to 450k methylation levels, we used data from the same samples using the two methods, and compared their methylation levels at the same sites both graphically (Figure 1D) and assessing the mean differences and root-mean-square deviation (Figure 1C). We then validated our model’s findings by applying the same model to the microarray data to overlapping sites and considered loci significant at FDR < 0.05 and validated if it’s association had p < 0.05 in the validating set (Figure S3).

**Figure 1:**
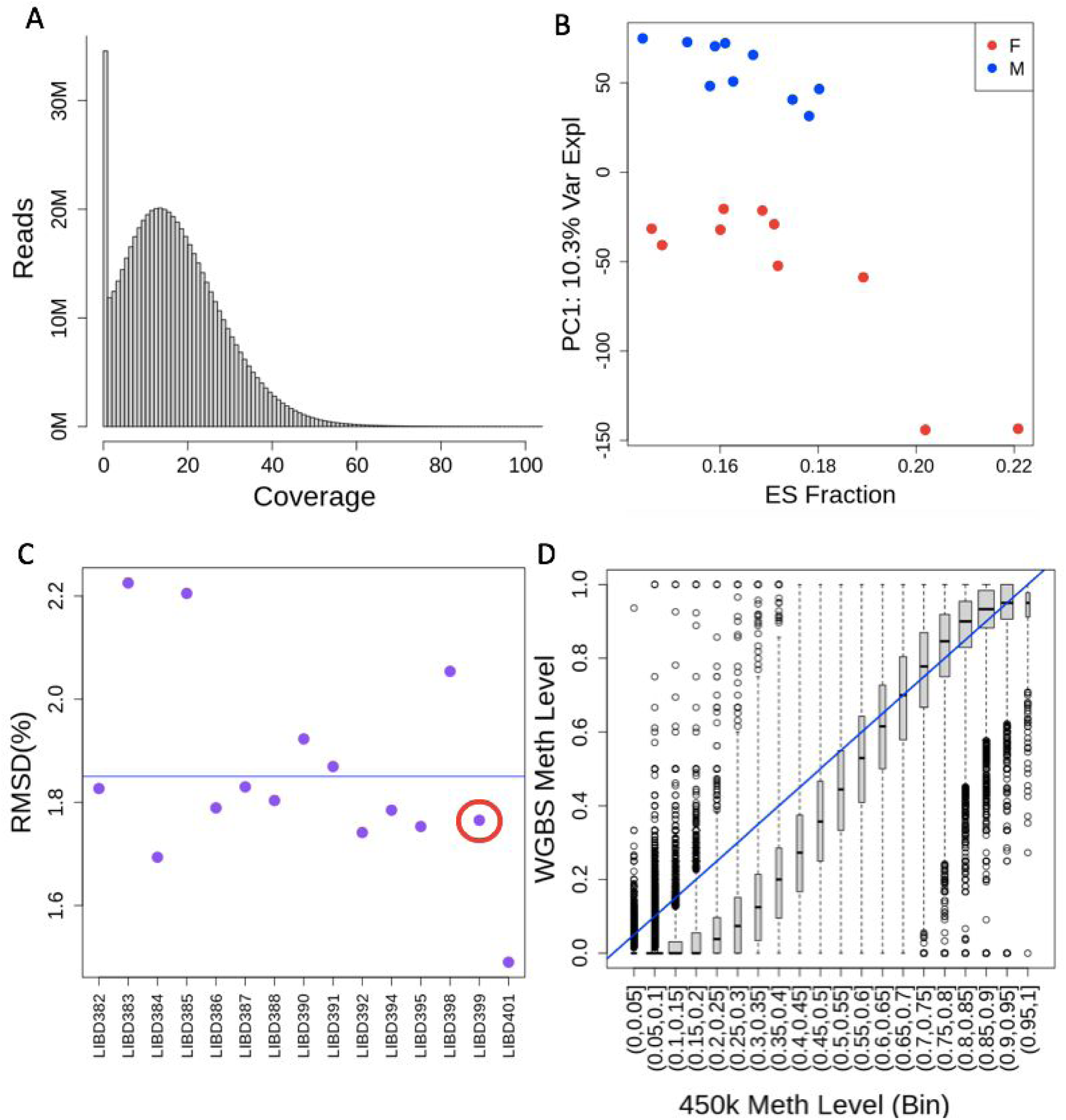
WGBS Data. **(A)** Coverage levels across all CpGs. The mean coverage level was 17.7 reads per CpG. **(B)** PCA on all CpG DNAm levels shows that the top principle component – accounting for 10.3% of variance, is sex. However we can also see that there is another factor separating the two clusters of females, shown here to be embryonic stem (ES) cell fraction. **(C)** Root-mean-square deviation (RMSD) of WGBS measurements from 450k measurements for each sample. Mean RMSD is shown. **(D)** WGBS measured DNAm for each site on 450k, organized by 450k methylation measurement, of sample LIBD399, show in the red circle in (C).

### Gene set enrichment

We annotated our data using Gencode v. 29 on hg38. We performed gene ontology and gene set enrichment using clusterProfiler (v3.12)^18^ with a p-value cutoff of 0.01 and q-value cutoff of 0.04. We used SFARI 2.0 ^19^ for an autism spectrum disorder gene set, and a set of clinical gene sets defined by Birnbaum et al for other neuropsychiatric and neurodevelopmental disorders ^20^. Enrichment was calculated on a background of genes expressed in our samples, to avoid brain bias. We performed LD score regression as described by and with data from Finucane et al ^21^.

### Data Analysis

Cell composition of samples was deconvoluted using flow-sorted DLPFC microarray data from fluorescence-activated nuclear sorted neurons and non-neurons ^22^ and R package minfi ^23^. Linear regression modelling was performed accounting for sex, age, and embryonic stem cell content. Sex-related CpGs and their surrounding sequence were re-aligned to the genome with Bowtie2 to check for homology. Association was tested using limma (v3.30) ^24,25^ to create a linear regression model accounting for sex, age, and embryonic stem cell content.

### RNA-seq comparison analysis

The homogenate RNA-seq samples were also part of a larger study of RNA-seq data from homogenate DLPFC tissue (BrainSeq Consortium Phase I) ^26^. We compared a site’s methylation or mean methylation (for DMRs) to the expression level from RNA-seq using a linear model adjusting for the strongest covariates, including sex, age, and estimated embryonic stem cell (ESC) content.

## Results

### Section 1 Whole genome bisulfite sequencing in the prenatal human cortex

We performed whole genome bisulfite sequencing (WGBS) to better characterize the shifting DNAm landscape in the developing human dorsolateral prefrontal cortex (DLPFC) in 20 prenatal samples during the second trimester *in utero* (Table S1). After data processing and quality control (see Methods), we analyzed 28,612,402 CpG and 13,166,520 CpH (H=A, T, or C) sites across the epigenome. We first focused on CpG DNAm and performed a series of quality control analyses. We quantified read coverage at every CpG, which after processing and alignment resulted in an average of 17.7 reads per CpG (Figure 1A). Most of these sites are methylated (>80% DNAm) and a minority are unmethylated (<20% DNAm) (Table S2). We then performed principal component analysis (PCA) on the DNAm levels across these high-coverage CpGs and found that the top principal components were most associated with sex and estimated fractions of embryonic stem cells (ESCs) via deconvolution (Figure 1B; Figure S1; see Methods), in line with our previous work ^9^. We lastly compared the DNAm levels from WGBS to levels from Illumina 450k microarray measured on 15 of the 20 samples for the subset of CpGs in common to both technologies (N=456305), and found that overall, CpG DNAm methylation levels were highly concordant regardless of CpG read coverage (Figure 1C-D). These analyses together suggested that our DNAm data was of high quality and available for subsequent differential methylation analysis.

### Section 2 DNAm associations and their characterization

To understand the changes in the prenatal epigenetic landscape, we performed linear modeling across all CpG sites. We found that DNAm changes were abundant even during this relatively restricted period in prenatal development, with 36,546 CpG sites differentially methylated across the ages of 14-20 post-conception weeks (PCW, at FDR < 0.05, Table S3). On average, each week of development was associated with a 1.8% change in DNAm (IQR: 1.18% - 2.33%) at a given CpG, with some sites changing as much as 8.7% per week. These differentially methylated CpGs were not evenly distributed across the genome, with none on the Y chromosome, and a small number on the X chromosome (Table S4). There was also some unevenness on the autosomes, with some chromosomes having up to 2.5x higher frequency of age-associated CpGs than others (Table S4). 2,743 of these CpGs lie in previously annotated CpG islands, with 5,933 in shores, defined as 2 kb from island ends. The vast majority (73%) of these CpGs lie within genes, and 2,932 lie within 1 kb of a transcription start site and thus potentially in promoters. This leaves 6.5% in intergenic space. Sites were fairly evenly split in the direction of methylation change by age, with 46% increasing in methylation as the cortex develops. Additionally, less than half of these CpG sites are significantly associated with the estimated ESC proportion in the sample, suggesting that many of these CpG sites have true prenatal age changes in methylation rather than reflecting effects of maturing cell type composition.

We further explored whether these sites could be organized into differentially methylated regions (DMRs), which have been shown to be more functionally relevant than individual CpGs. We therefore used a “bumphunting” technique adapted to WGBS data ^17^ to identify regions of methylation with 1.5% DNAm changes per week (corresponding to 10.5% changes across our developmental window) change in DNAm levels across adjacent CpGs, and calculated statistical significance using 1000 bootstrap-based permutations ^27^. Using a conservative cutoff (FWER < 0.2) there were 34 DMRs across prenatal development, though a less conservative cutoff (p < 0.05) identified 3,446 DMRs (Table S5, Figure S2A for DMR plot). The DMRs are similarly unevenly distributed throughout the genome as the CpGs, being far less frequent on chrX and variable among the autosomes. The DMRs on average had a width of 1696 bp (IQR: 921-2181), 75% overlap with annotated genes, and 12% overlap with annotated CpG islands. Like the CpGs, the DMRs were split between hypo- and hypermethylation, where in 49% of the DMRs, methylation increased with age.

We also found differentially methylated sites and regions between the sexes in these prenatal samples. There were 618,978 significantly differential CpGs by sex and as expected, the vast majority (95%) of these were on the sex chromosomes. There were still 30,823 significant autosomal CpGs (Table S6), and while 993 of them were in regions homologous to the sex chromosomes, the majority (97%) had no homology to chrX or chrY. Among these, we found conservatively 28 DMRs with a 10% methylation change between sexes (Table S7, Figure S2B for DMR plot). Again we saw that these were not global, but regional changes in methylation, with equal numbers of DMRs being hypo- and hypermethylated in males and females. These data show that there are many differences in the DNAm landscape throughout second trimester development, and between sexes prenatally.

Previously, many studies of brain DNA methylation have used the Illumina Infinium® HumanMethylation450 BeadChip (“450k”) and more recent Infinium MethylationEPIC (“850k”) microarray technologies. While these platforms can sensitively measure DNAm levels without the high coverage/sequencing requirements of WGBS, they assay a limited number of sites. To better identify the tradeoff between breadth (assaying more sites) and depth (assaying a given site more accurately), we compared our WGBS findings to analogous statistics calculated using these arrays on the same DNA extractions. First, using the probe coordinates alone, the 450k array does not measure DNAm levels at 35,401 (97%) of the significant age-correlated CpGs or 610,578 (98.7%) of the significant sex-correlated CpGs we identified. The 450k DNAm data validated 566 (49%) out of the 1,145 age-correlated sites that were covered. Performing the same age model on the 450k data, our WGBS data validated 279/423 (66%) of the significant results. The effects of age on methylation were generally directionally concordant between the two datasets (Figure S3). Though DMRs have much wider spans, the 450k does not cover 10 (30%) and 12 (43%) of DMRs for age and sex, respectively. The newer 850k microarray had almost twice the number of probes, yet we found that it does not measure 95% of our significant sites for age and 98% of sites for sex. Additionally, it still failed to capture 24% of age DMRs and 36% of sex DMRs. Microarrays are potentially missing a great deal of significantly differential sites, suggesting WGBS is best for thorough analysis.

### Section 3 Functional characterization of differential sites

While differential methylation analysis provides detailed information about changes in the methylation landscape across brain prenatal development and between the sexes, we wanted to better understand the molecular consequences of these changes. We performed gene set enrichment and gene ontology to better understand in which processes the genes containing significant CpGs are involved. The top biological processes associated with genes containing differentially methylated CpGs across age were related to axon development and guidance, and regulation of neuron projection (Figure S4, Table S8). These sites were also shown to be enriched for enhancers in fetal brain tissue as well as enriched for transcription start site regions in brain tissue from the Epigenome Roadmap project ^28^, implying that these areas are likely functional in prenatal brain ^28^ (Figure 3A). To better assess the putative functionality of the methylation at these CpGs and DMRs, we correlated these DNAm associations with nearby expression levels using RNA-seq data from the same cortical dissection. Among the 36,546 age-related CpGs, 59% were within an RNA-seq-measured gene (corresponding to 1,630 unique genes). 7,998 (37%) of these CpGs had methylation levels significantly correlated (p < 0.05) to expression of the gene they lie directly within (661 unique genes, 41%, see Methods, Figure 3B-C). These expression-correlated CpGs did not have any differential gene ontology from the overall group of genes. These sites were mostly in weak transcription, enhancer, and quiescent chromatin states in fetal brain (Figure S5) ^28^. With an expanded window of 5 kb around genes, 24,679 (66%) CpGs were accounted for, and 8,759 (35%) correlated to the nearest gene’s expression. It is important to note that this only tested sites within actual genes, and only the effect on the same gene, so enhancer, upstream, and trans-acting effects were not accounted for.

**Figure 2:**
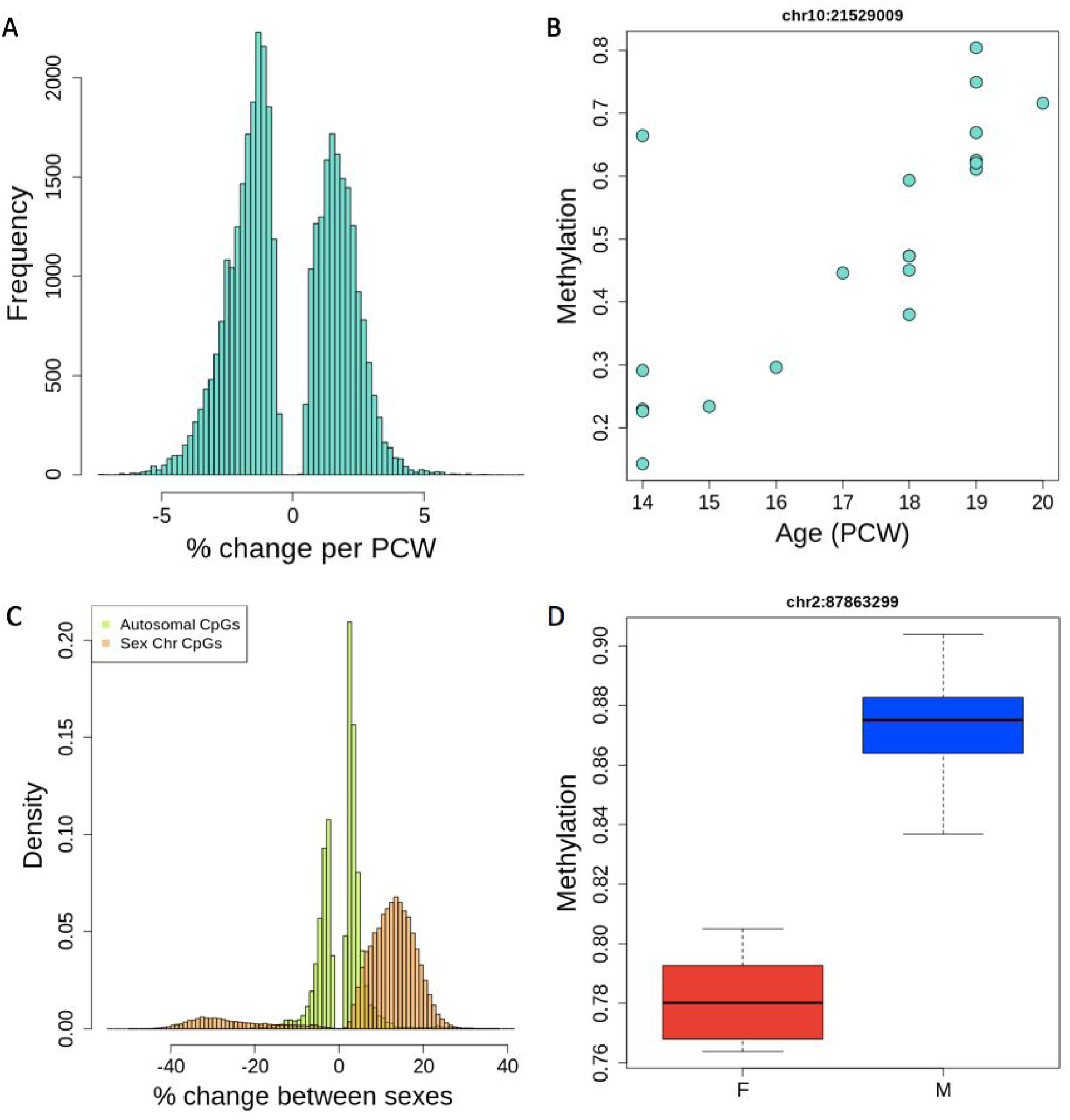
Differential methylation across ages and between sexes. **(A)** Percent change in methylation per post conception week (PCW) across all significantly age-differential CpGs (FDR < 0.05). **(B)** An example of an age-differentially methylated CpG on chromosome 10. **(C)** Percent change in methylation between sexes at sex-differential CpGs, showing that sex chromosomes tend to have greater percentage differences than autosomal CpGs. **(D)** An example of a sex-differentially methylated CpG on chromosome 2.

**Figure 3:**
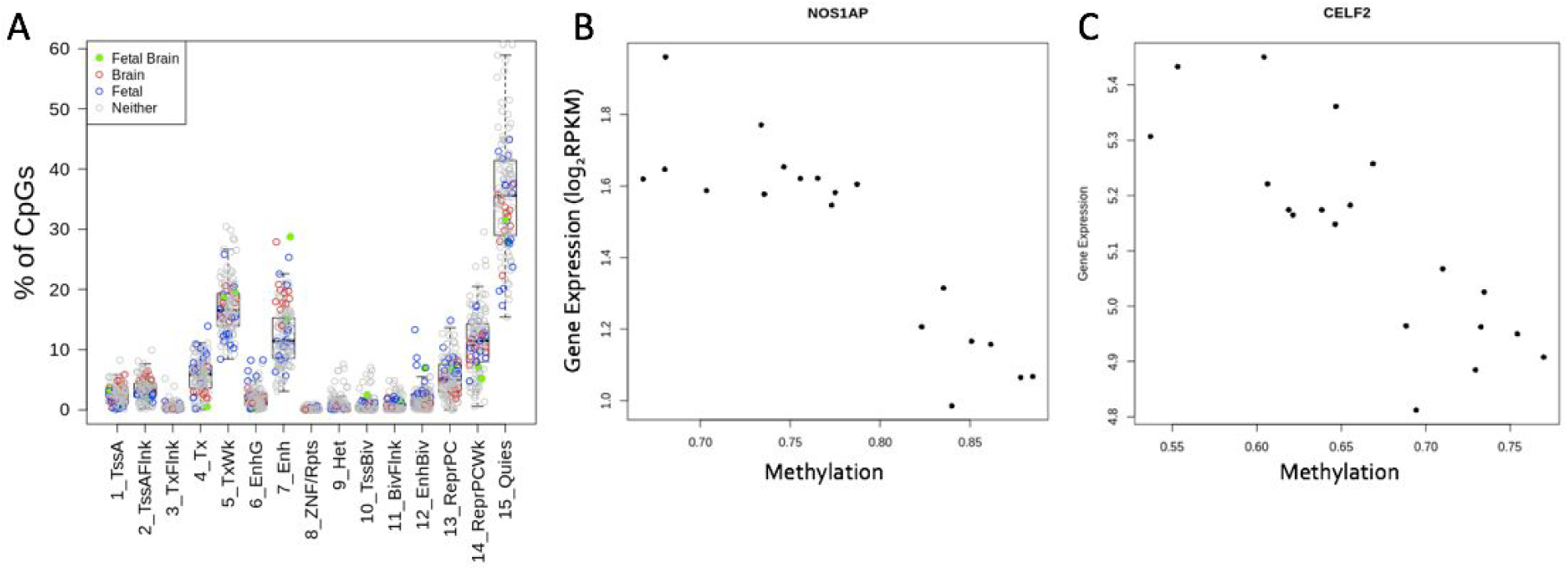
Functional consequences of differential methylation. **(A)** Chromatin states of age-associated CpGs in various tissues. We show the distribution of chromatin states from the 15-state model of our CpGs in various tissues from the Roadmap Epigenomics Consortium. In Roadmap fetal brain tissues, our sites are enriched for enhancer regions, and in all brain tissues, our sites are enriched for TSS sites. **(B, C)** Examples of correlation between methylation level and gene expression. Gene expression from RNA-seq is plotted against DNAm level at correlated sites, showing a decrease in gene expression with increasing methylation in two example genes.

The correlation to expression was much lower for sex-related CpGs, at 5% and 9% significant correlation depending on inclusion or exclusion of sex chromosomes. The lack of correlation to expression here is unsurprising because there are almost no transcriptomic differences between the sexes in these samples. Gene set enrichment on autosomal sex-related CpGs found genes involved with synaptic transmission and signaling, regulation of GTPase activity, and the glutamate receptor signaling pathway (Table S9).

Age-related DMRs were most enriched for genes related to stem cell proliferation and various cell fate specifications (Table S10). Among the DMRs across age, 21% were significantly correlated with gene expression. The genes represented by this are *OTX1, AC246817.1, CYP2E1, PLEKHH3*, and *DUX4L32*. There was no enrichment for any gene category among the sex-related DMRs, but it is worth noting that many of these genes coded for lncRNAs. 25% of sex-related DMRs were correlated to gene expression within their own gene and these genes were *LINC01606, LINC01166, SPATC1L*, and *RFPL2*. Correlation to gene expression overall was not different between differentially methylated regions and non-differentially methylated regions; the DMRs we found were not enriched for highly correlated regions. These results provide potential starting points for further understanding the DNA methylation landscape of the developing brain, allowing us to understand which processes are active during normal and disordered development.

### Section 4 CpH methylation changes

While DNA methylation occurs exclusively in the CpG context in most somatic cell types, neurons in the human brain uniquely have high levels of DNA methylation in other cytosine contexts (predominantly CpA) ^29^. We therefore investigated the potential role of CpH methylation across brain development. Only 2.4% (IQR: 2.33-2.49) of sites in a CHG context and 1.71% (IQR: 1.61-1.79) of sites in a CHH context were methylated. For comparison, around 10% of CpH sites are methylated in postnatal neurons, so it is likely as previously suggested ^1^ that CpH methylation accumulates beginning around the time of birth. This contrasted with CpG sites, which were predominantly methylated across the genome (Table S2). Because there were so many more CpH than CpG sites in the genome, this means that there are actually similar numbers of methylated CpG and CpH sites, despite the different proportions. Using the same linear model as was used for CpGs on 13,166,520 CpH sites, no CpHs were significantly associated with age within this time period when accounting for the large multiple testing burden. However, 51 autosomal CpHs were significantly associated with sex (at FDR < 0.05, Table S11). Additionally, there were 40 associated CpHs on the sex chromosomes, though most of these were on chrY, overall very proportionally different from CpG methylation. There was no enrichment for any more specific trinucleotide context in these significant sites over the whole genome. These CpHs were less likely to be in or near genes than the CpGs, but of the 17 autosomal CpHs that were in or near genes, only 1 was significantly correlated (p < 0.05) with expression of the nearest gene within 10 kb. It is notable that the majority of these CpHs are relatively far from a gene. Additionally, the effect size of the CpH methylation at these significant sites is independent of the effect of CpGm in the surrounding area in most cases. Five of the CpHs’ effect is reduced when accounting for the methylation of the nearest CpG (Figure S6), showing that these few are dependent on the local CpG landscape, but most function independently. These loci are found both in and outside of genes and their surrounding regions. Overall, CpH methylation does not seem to be very dynamic or functional in prenatal second trimester development, though later in early postnatal life it is ^30^.

### Section 5 Links to neuropsychiatric conditions and their (genetic) risk

To understand the clinical implications of our findings, we tested the genes represented by prenatal age-associated CpGs and found that they were enriched for bipolar disorder- (p = 0.035), neurodevelopmental disorder- (p = 0.0011), and schizophrenia- (p = 0.00022) associated genes ^20^. Age-differential CpGs with correlation to gene expression were not further enriched. The set of age-differential CpG genes was particularly enriched for autism-associated genes (p = 9×10^−20^, OR = 5.6), with 163 of our 1,630 genes being linked to autism spectrum disorder (ASD) ^19^. Sex-differential CpGs were also enriched for ASD-associated genes (p = 6×10^−29^, OR = 2.9), but not for genes associated with other psychiatric disorders.

DMRs represented a greater portion of the genome than CpGs, and thus may have different functional effects. To further understand what phenotypes may be linked to our DMRs, we performed stratified LD score regression ^21^. Our DMR sets represented a fairly small portion of the genome, but at 5.8 Mb, the less stringent age-based DMRs were wide enough to detect enrichment of BMI- and Subjective Well-Being- (SWB) associated markers (Table S12). These DMRs were also shown to be enriched for brain-linked traits overall, compared to non-brain-linked traits. Additionally, the DMRs overlap with 15 of 108 schizophrenia loci from Psychiatric Genetics Consortium 1 ^31^. These results suggest that dynamic methylation states in prenatal development may play a role in many disease and non-disease phenotypes that manifest later in life.

## Discussion

Here we demonstrate that CpG methylation changes are associated with prenatal cortical development, are likely functionally relevant, and may play a role in developmental psychiatric disorders such as autism spectrum disorder (ASD).

The shifting DNAm landscape during this restricted prenatal time period - the second trimester - likely plays a role in neurogenesis of the developing human brain. Following current dogma, changes in DNAm at these critical sites presumably lead to altered gene expression which promotes the formation of new neural connections in the cortex. At this point in time, we see that it is CpGs that are the important sites in the DNAm landscape, as opposed to CpHs. This is in line with previous research which finds that most CpHm is established around birth and postnatally ^1,32^, later in neuron differentiation and development ^33^. We also see that much of this regulation is occuring in *cis* -within the site’s own gene. We do not currently understand the *trans* effects of our developmentally dynamic sites, but they may explain the effects of the methylation that does not correlate to its nearest gene’s expression. The fact that the age-associated CpG sites we identified are often in enhancer regions in prenatal brain is further evidence that these sites may be regulating from a distance.

Our findings are in line with previous findings based on a microarray platform^10^, but using WGBS allows us to assess methylation at far more loci than the more commonly used Illumina microarrays. We’ve shown that these microarrays do not even measure the majority of sites that are dynamic in neural development. Despite potentially less precision, we also show that WGBS has enough coverage to assess DNAm dynamics and that its findings were consistent with deeper-coverage assays. Additionally, we present novel loci due to our narrower developmental age range, which means that we can detect more subtle changes in the second trimester but also means that we do not replicate all the findings of studies done across wider timespans. Confining our samples to just cortex tissue rather than whole brain tissue also allows us to detect more regionally specific changes. Most variation in our data likely comes from different cell types in the cortex, which we partially accounted for in our model by adjusting for ESC estimated proportions. We believe that most of the cell types in this area at this time are neurons, but there is likely a range of maturities leading to variation in methylation ^11^. By accounting for ESC fraction in our model, we believe that the age affects we find are truly a result of prenatal age.

We also find that there are autosomal differences in DNAm between the sexes, though by our assessment they seem less functionally significant. They may be acting more in *trans* which makes it more difficult to assess significance, but there are also just generally far fewer transcriptomic differences between sexes than between ages. Sex-differential DNAm later in life has been linked to psychiatric genes, which has been proposed as a mechanism for sex differences in psychiatric disorders ^34^, so perhaps the early changing DNAm is laying the groundwork for future effects. Like Xia et al ^34^, we found that ASD-associated genes are differentially methylated between the sexes, but this does not readily translate into transcriptomic differences.

Given the enrichment in our developmentally-associated sites for psychiatric-linked genes, these data support the notion of early neurodevelopmental components for disorders such as schizophrenia ^7,35,36^. Genes implicated in bipolar disorder, schizophrenia, and ASD are clearly dynamic at this time point in life, so dysregulation at this stage could lead to vulnerabilities to these disorders later in life. Our data also implies that BMI and subjective well being - non-disease traits - could be linked to neural development at this stage of life.

As more data is generated, particularly through genome-scale methods like WGBS, we will be able to establish normal ranges of DNAm at all ages, which will undoubtedly provide insight into molecular dynamics in this hard-to-study period and organ, as well as give clues to where deviation from the norm is important. Unfortunately, WGBS still does not measure hydroxymethylation, another chemical modification thought to be epigenetically important. Our findings are only the beginning of what may be found given the limited sample size of our study, but even here we reveal important processes. There is room for much more characterization of these epigenetic marks, but it is clear that they are worth understanding. DNA methylation serves as an exciting potential avenue to understand neural development and psychiatric disorders. There are clear and functional changes in the neuronal DNAm landscape over this important window of brain development and between sexes, and further investigation will help elucidate unknown mechanisms in the brain.

## Supplemental Data

Supplemental Data include 6 figures and 12 tables.

## Acknowledgements

The authors thank the UMB Brain Bank at the Department of Pediatrics in the University of Maryland School of Medicine for the tissue provided. This project was supported by The Lieber Institute for Brain Development and by NIH grants R21MH102791 and R01MH112751. Finally, we are indebted to the generosity of the families of the decedents, who donated the brain tissue used in these studies.

## Data Availability and Web Resources

Raw and processed nucleic acid sequencing data generated to support the findings of this study are part of the PsychENCODE Consortium and the Brainseq Consortium data releases. Specifically, WGBS data have been deposited at www.Synapse.org along with the other PsychENCODE data, under the accession code syn5842535. The homogenate RNA-seq samples were also part of a larger study of RNA-seq data from homogenate DLPFC tissue (BrainSeq Consortium Phase I) ^26^, which was also deposited at www.synapse.org and summarized in http://eqtl.brainseq.org/phase1. The processed, homogenate RNA-seq data for this study have additionally been deposited via Globus under the jhpce#brainepi-cellsorted collection at the following location: http://research.libd.org/globus/jhpce_brainepi-cellsorted/. NeuN-sorted RNA-seq data were originally published as part of phase II of the Brainseq Consortium (http://eqtl.brainseq.org/phase2/) and have also been deposited via Globus under the jhpce#brainepi-polyA collection at the following location: http://research.libd.org/globus/jhpce_brainepi-polyA/. Publicly available data reprocessed in support of the conclusions in this work were downloaded from the Gene Expression Omnibus under GEO accession GSE47966.

