## Supplemental Figures and Table Legends for "Characterizing the dynamic and functional DNA methylation landscape in the developing human cortex"

**Supplementary Figures and Table Legends**

**Supplementary Figures**

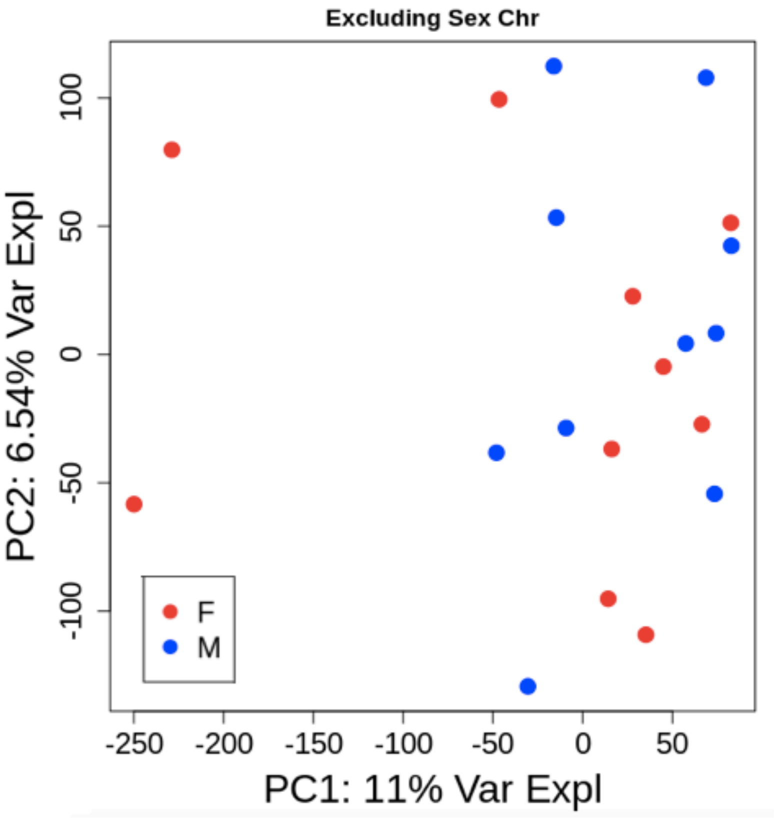


**Figure S1**: **PCA excluding sex chromosomes.** Principle components of variance in DNAm levels. Performing PCA on the data without the sex chromosomes, we see that the sex differentiation falls out, and ESC fraction becomes the primary component.

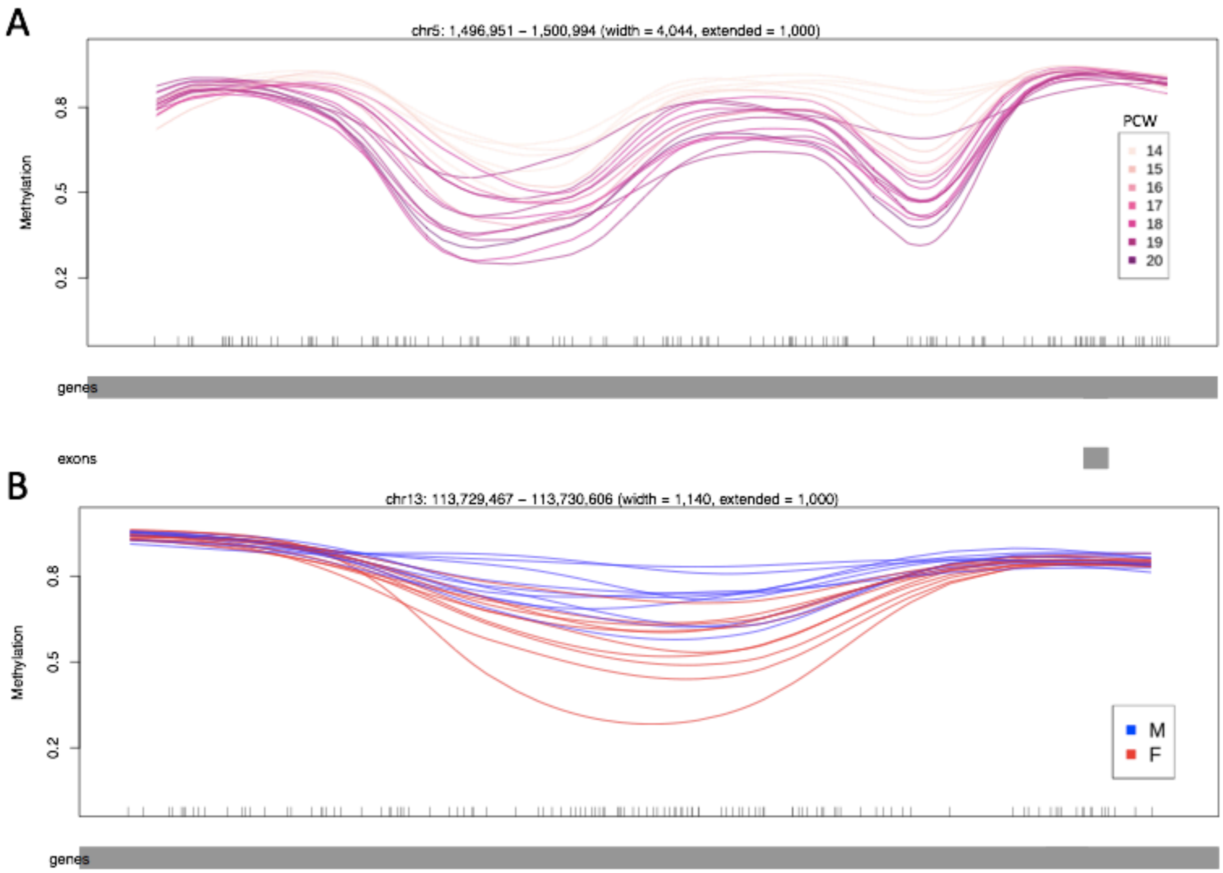


**Figure S2: DMRs. (A)** a DMR in *LPCAT1*, in which DNAm decreases with age. **(B)** a DMR in *GRK1*, in which males have higher DNAm

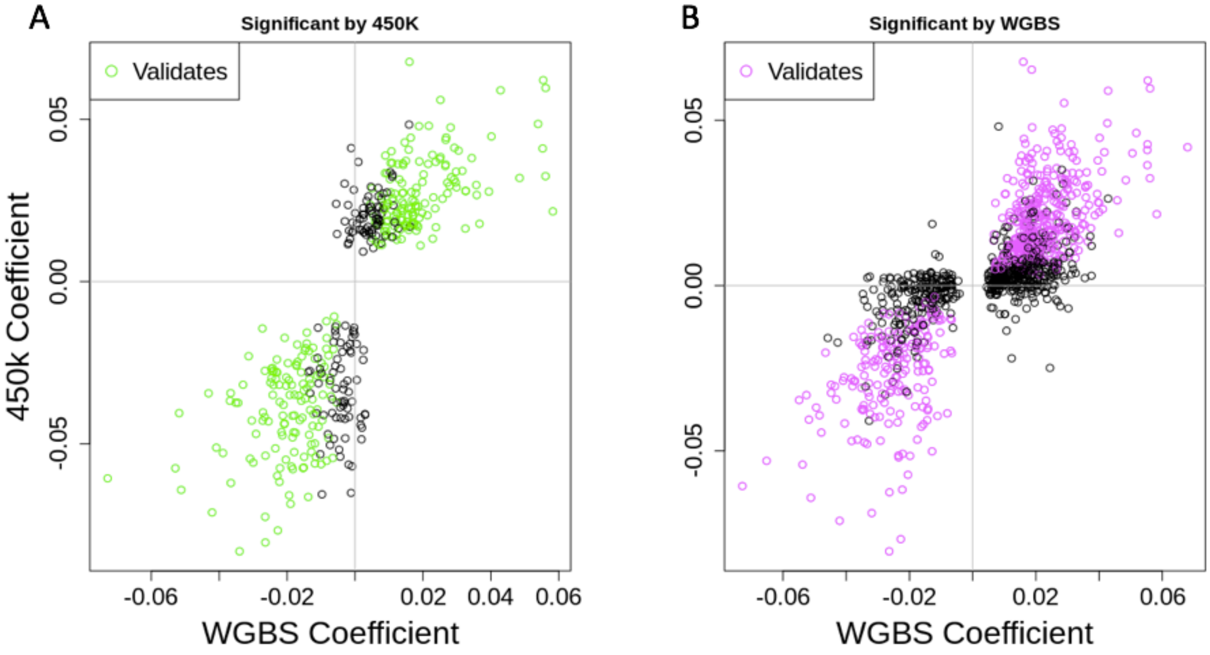

**Figure S3: Validation of findings between 450k and WGBS**. **(A)** We perform the same linear regression modelling on data from the 450k microarray to see which sites it identifies as significantly age-associated. We plot the effect coefficient by that model against the effect coefficient for the same sites in the WGBS model. Sites are considered validated if the coefficients are directionally concordant and the p-value from the WGBS model is < 0.05. **(B)** A reverse of (A), we take points that are significant by the WGBS model, then check if they validate by the 450k model. Overall we see great directional concordance between the two models.

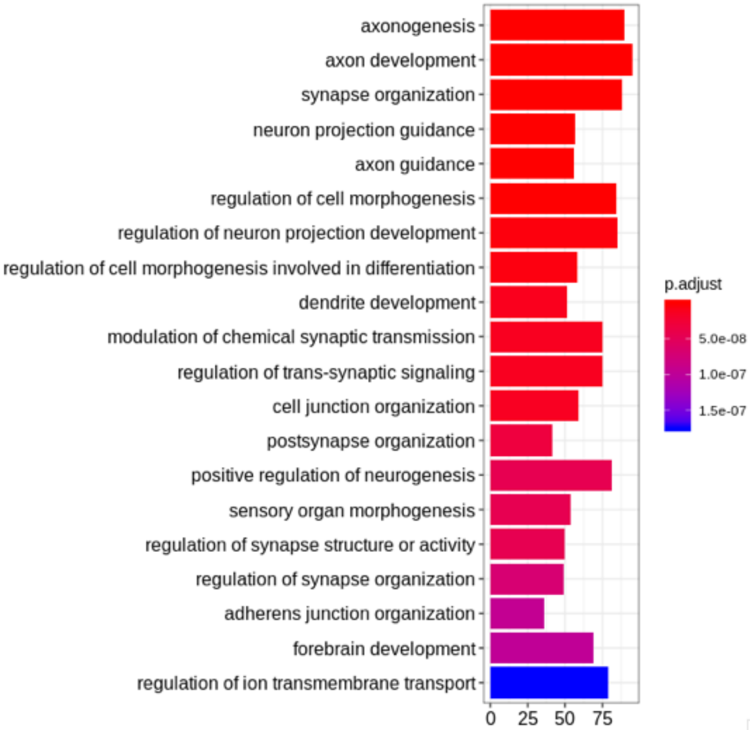


**Figure S4: Gene Ontology and gene set enrichment for age-associated CpGs**. The top 20 GO hits for all categories are plotted in order of significance (indicated by bar color), showing the gene count for each GO category.

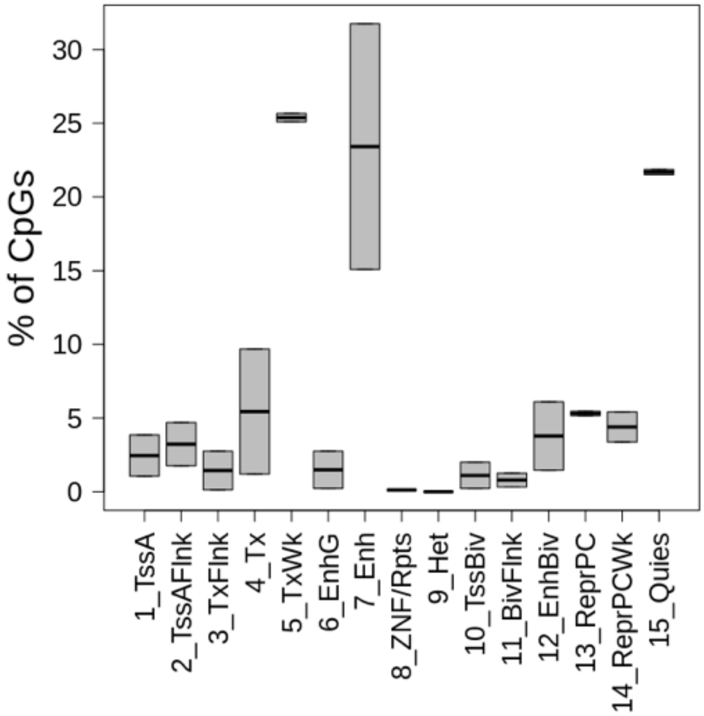


**Figure S5: Chromatin state of CpGs correlated to expression in fetal brain tissue.** We see that in fetal brain tissue, most of our age-associated and expression-correlated CpGs are in weak transcription or enhancer states.

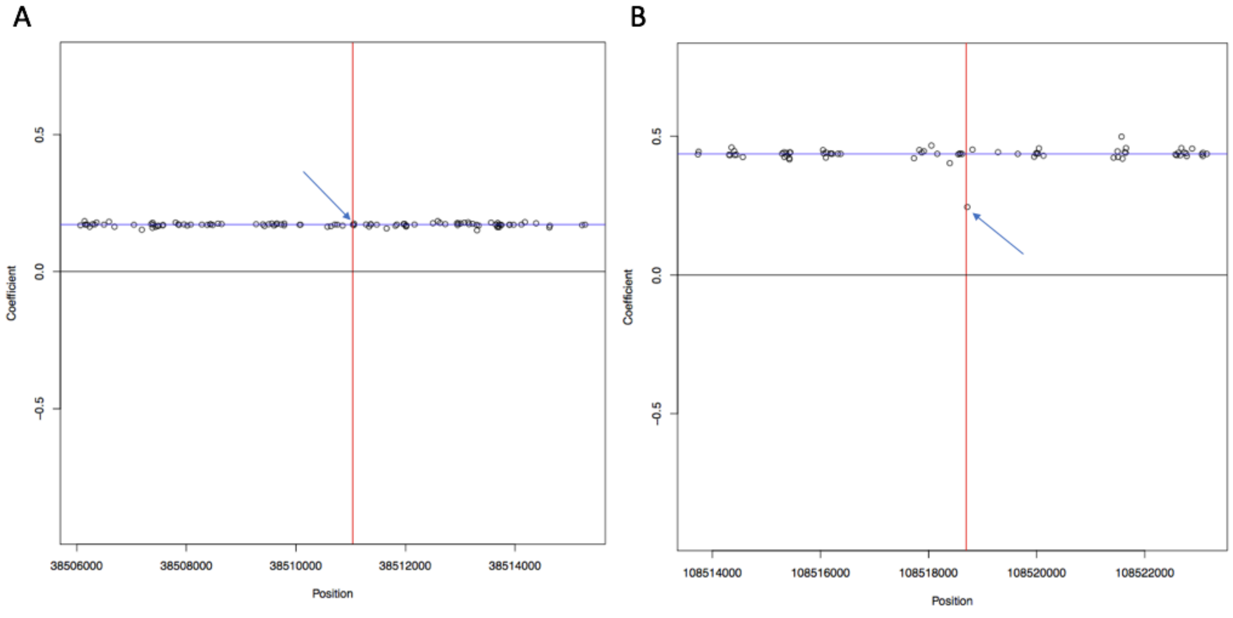


**Figure S6: CpH effect independence of nearby CpGs**. **(A)** A representative plot of CpH effect size (blue line) at a CpH site (red line), plotted against CpH effect size when CpGm is included in modelling (points). Most sites look like this. **(B)** 5 CpHs have reduced effect size when accounting for CpGm of the nearest CpG, shown in this example.

**Supplementary Table Legends – Supplementary Tables in separate sheets of a separately submitted Excel file**

**Table S1: Sample phenotype data**
MacrogenID & brnum: sample identifiers
ageweeks: age in weeks post conception
sex: F = female, M = male
race: AA = African American, CAUC = Caucasian
DA_NEURON: estimated dopaminergic neuron fraction
ES: estimated embryonic stem cell fraction
NeuN_neg: estimated NeuN – cell fraction
NeuN_pos: estimated NeuN + cell fraction
NPC: estimated neural progenitor cell fraction

**Table S2: Methylation State of CpGs**

**Table S3: Age-associated CpGs**
seqnames: chromosome
start and end: hg38 position
nearestgene: nearest gene
CorrelatedtoExpression: correlated to expression of the nearest gene ($p < 0.05$)
change: change in methylation per week. Positive values indicate increasing M with age
adj.p.val: FDR corrected p value

**Table S4: Chromosomal Distribution of age-associated CpGs**

**Table S5: Age differentially methylated regions**

**Table S6: Sex-associated CpGs**
seqnames: chromosome
start and end: hg38 position
nearestgene: nearest gene
CorrelatedtoExpression: correlated to expression of the nearest gene ($p < 0.05$)
change: change in methylation between sexes. Positive values indicate hypermethylation in males
adj.p.val: FDR corrected p value

**Table S7: Sex differentially methylated regions**

**Table S8: Age-associated CpG Gene Ontology**
ONTOLOGY: Gene Ontology type: CC = cell compartment, BP = biological process, MF = molecular function
ID: Gene Ontology ID
Description: Gene Ontology set description
GeneRatio: Fraction of differentially expressed genes were in the GO set
BgRatio: Fraction of differentially expressed genes that were not in the GO set
Pvalue: P-value resulting from hypergeometric test
p.adjust: Benjamini-Hochberg adjusted p-value (FDR)
qvalue: Storey adjusted p-value

geneID: Gene symbols corresponding to the differentially expressed genes in the GO set (ie from GeneRatio above)
Count: Number of differentially expressed genes in the GO set (numerator of GeneRatio, to avoid forced Excel conversion to dates from some fraction)

**Table S9: Sex-associated CpG Gene Ontology**

**Table S10: Age DMR Gene Ontology**

**Table S11: Significantly sex-associated CpHs**

**Table S12: LDSC statistics for age-associated DMRs**
